# Oral Health-Related Quality of Life in Children and Adolescents with Osteogenesis Imperfecta: cross-sectional study

**DOI:** 10.1101/424812

**Authors:** Mohammadamin Najirad, Mang Shin Ma, Frank Rauch, Vernon Reid Sutton, Brendan Lee, Jean-Marc Retrouvey, Members of the BBD, Shahrokh Esfandiari

## Abstract

**Background:** Osteogenesis imperfecta (OI) affects dental and craniofacial development and may therefore impair Oral Health-Related Quality of Life (OHRQoL). However, little is known about OHRQoL in children and adolescents with OI. The aim of this study was to explore the influence of OI severity on oral health-related quality of life in children and adolescents.

**Methods:** Children and adolescents aged 8-14 years were recruited in the context of a multicenter longitudinal study (Brittle Bone Disease Consortium) that enrolls individuals with OI in 10 centers across North America. OHRQoL was assessed using the Child Perceptions Questionnaire (CPQ) versions for 8 to 10-year-olds (CPQ_8-10_) and for 11 to 14-year-olds (CPQ_11-14_).

**Results:** A total of 138 children and adolescents (62% girls) diagnosed with OI types I, III, IV, V and VI (n=65, 30, 37, 4 and 2, respectively) participated in the study. CPQ_8-10_ scores were similar between OI types in children aged 8 to 10 years. In the 11 to 14-year-old group, CPQ_11-14_-scores were significantly higher (i.e. worse) for OI types III (24.7 [SD 12.5]) and IV (23.1 [SD 14.8]) than for OI type I (16.5 [SD 12.8]) (P<0.05). The difference between OI types was due to the association between OI types and the functional limitations domain, as OI types III and IV were associated with significantly higher grade of functional limitations compared to OI type I.

**Conclusion:** The severity of OI impacts OHRQoL in adolescents aged 11 to 14 years, but not in children age 8 to 10 years.

## Background

Osteogenesis Imperfecta (OI), also known as “brittle bone disease,” is a rare heritable disorder (prevalence 8 per 100,000 people) that is characterized by recurrent fractures and, in severe cases, skeletal deformities [1]. Extra-skeletal features such as blue or grey discoloration of sclera and discoloration and brittleness of teeth (dentinogenesis imperfecta, DI) can be associated. In about 90% of individuals with a clinical diagnosis of OI, a dominant mutation in the genes that code for type 1 collagen alpha chains (*COL1A1* and *COL1A2*) can be identified as the cause of the disease [2]. OI has traditionally been classified into four clinical types reflecting the severity of the phenotype (type I – mild; type II - neonatal lethal; type III – severe; type IV - moderately severe). The number of OI types has subsequently been expanded based on distinct clinical features and later based on the underlying genetic etiology [1, 3]. There is presently no cure for the disease, but pharmacological therapies using bisphosphonate drugs are widely used to strengthen bones, decrease the pain and fracture rates [3].

The World Health Organization (WHO) defines quality of life (QoL) as the “individual’s perception of their position in life in the context of the culture and value systems in which they live and in relation to their goals, expectations, standards, and concerns” [4]. The term “Health-Related Quality of Life” (HRQoL) narrows QoL to aspects relevant to health [5], and the term “oral health-related quality of life” (OHRQoL) focuses on physical, psychological, and social impacts of oral and orofacial conditions and disparities in oral health on overall health and QoL of individuals.

The pathological effects of OI on dental tissues and oral cavity usually develop in early life and may therefore influence OHRQoL during childhood and adolescence. Orofacial manifestations often associated with OI include DI, posterior open bite (lateral open bite), class III dental and skeletal malocclusion, anterior and posterior crossbites and impacted teeth [6]. The degree of the oral manifestations seem to be most severe in OI type III, as this type is associated with more severe craniofacial deformities and a higher prevalence of DI than milder forms of OI, such as OI type I and IV [7, 8]. Until now little is known about OHRQoL in children with OI. It is unclear whether OHRQoL differs between OI types and between children and teenagers with OI.

In the present study we therefore assessed OHRQoL in children and adolescents with OI who participated in a multicenter study exploring the natural history of the disease. We hypothesized that the more severe phenotype in OI type III would result in lower OHRQoL compared to less severe OI types.

## Materials and methods

### Study participants, recruitment, and setting

Study participants were recruited through the Brittle Bone Disease Consortium [9] that comprises several specialized centers from across North America (Houston, Montreal, Chicago, Baltimore, Portland, Washington DC, New York, Omaha, Los Angeles, Tampa). The consortium is a Rare Disease Clinical Research Network that is funded by the National Institutes of Health. One of the projects conducted by the consortium is a natural history study. The study was approved at all participating study centers, and all study participants or their legal guardians provided informed consent.

The present evaluation includes all children and adolescents with any OI type for whom OHRQoL data could be obtained in the first two study years from 6^th^ of August 2015 to 3^rd^ of August 2017. As the two pediatric OHRQoL instruments used in the study were specific for the age ranges from 8 to 10 years and from 11 to 14 years, respectively, the present analysis includes children and adolescents from 8 to 14 years of age. OHRQoL questionnaires were collected on paper at participating study sites and entered into an online data capture system that is maintained by the study Data Management and Coordinating Center (University of South Florida).

### Data collection

OHRQoL was evaluated using the Child Perception Questionnaire (CPQ). The CPQ_8-10_ containing 25 questions was used for children between 8 and 10 years of age [10], the CPQ_11-14_ comprising 37 questions was used for individuals aged 11 to 14 years [11]. Study participants were asked to complete the questionnaire unassisted by parents or investigators [12, 13]. These instruments are comprised of four health domains: oral symptoms, functional limitation, emotional well-being and social well-being related to oral health conditions. All questions consider the frequency of events in relation to the condition of the mouth or teeth over the previous four weeks (CPQ_8-10_) or three months (CPQ_11-14_). The responses to questions are scored on a frequency scale using the following response options and associated codes: ‘Never = 0’; ‘Once/twice = 1’; ‘Sometimes = 2’; Often = 3’, and ‘Everyday/Almost every day = 4’. The questionnaires also contain two single-item global ratings. Additive subscale CPQ scores (domain-specific score) are computed by summing response codes. The overall CPQ scores are computed by adding up all four domain subscale scores, which may range from 0 to 100 for CPQ_8–10_ and 0 to 148 for CPQ_11–14_. Higher scores denote worse OHRQoL [10-12, 14]. The validity, reliability, and responsiveness of this measure have been established in various settings [15-19].

### Data analysis

Statistical analyses were performed using Stata 13.0 software (StataCorp. 2013. *Stata Statistical Software: Release 13*. College Station, TX: StataCorp LP) with a 5% significance level. Collected variables were classified into three levels: (1) Sociodemographic characteristics; (2) medical and physical conditions; and (3) OHRQoL. Missing values for some CPQ constituent questions (6% of questionnaire data fields) and were substituted with the mean value of that variable across each OI type (single imputation method). Descriptive and univariate analyses were performed across different types of OI separately for each age group. Welch’s t-test (independent samples t-test) was employed to handle the unequal variances and sample sizes between the groups of binary variables. When the sample size of a group was less than 15 patients, the Mann-Whitney U-test (non-parametric) was performed to assess for the significance of differences between two groups. To determine the significant relationship between categorical variables, Chi-square test or Fisher’s exact test for contingency tables with small cell counts were employed. CPQ scores and their constituent subscale scores were transformed to ordinal variables using their 33^rd^ and 66^th^ percentiles. Age, gender, race, and having a family history were identified as the minimum set of potential confounders, to be included in the multivariate analyses. Multivariable ordinal logistic regression analyses were employed to estimate the total effect of OI types on CPQ score and its constituent domains.

## Results

A total of 138 individuals (62% females) aged 8-14 years (11.6 ± 2.1 years) and diagnosed with OI types I, III, IV, V and VI (n=65, 30, 34, 6 and 3, respectively) participated in the study. As the number of participants with OI types V and VI was too small for meaningful statistical analysis, these two OI types were analyzed in one group (Others) (Table 1). All patients were living with their biological parents. Six participants (OI type I, n=3; OI type III, n=1; OI type IV, n=2) were homeschooled, the others attended school.

In children aged 8 to 10 years (Table 1), more children with OI types I and IV were having a parent or ancestor living with OI compared to OI type III (p-value <0.05). All children with OI types III and IV had a history of bisphosphonate treatment (oral or IV) compared with 42% OI type I. Using wheelchair as a mean of transportation was more prevalent in OI type III compared to OI type I and IV (p-value <0.05). Number of days which the child has missed the school due to his/her condition related to OI was significantly more prevalent in type III in comparison to type I (p-value <0.05) and also significantly higher in type IV compared to type I (p-value <0.05). In the group of children (aged 8–10 years), there were no statistical differences in total scores of the CPQ8–10 or domain scores when different types of OI were compared (Table 2).

**Table 1.** Characteristics of the 8 to 10-year-old group

| Patients aged between 8-10 | OI I | OI III | OI IV | Others | All |
| --- | --- | --- | --- | --- | --- |
| **<u>Sociodemographic Characteristics</u>** |  |  |  |  |  |
| Enrolment number – n (%) | 26 (46) | 16 (29) | 11 (20) | 3 (5) | 56 (100) |
| Female | 13 (50) | 12 (75) | 7 (64) | 2 (66) | 34 (61) |
| Age – mean (SD) | 9.3 (1.0) | 9.2 (0.9) | 9.8 (0.5) | 9.6 (0.5) | 9.4 (0.9) |
| Race (White) – n (%) | 22 (85) | 13 (81) | 5 (45) <sup>c</sup> | 3 (100) | 43 (77) |
| others | 4 (15) | 3 (19) | 6 (55) <sup>c</sup> | 0 (0) | 13 (23) |
| Insurance status (Private) – n (%) | 20 (77) | 8 (50) | 8 (73) | 2 (67) | 38 (68) |
| Medicare/Medicaid | 6 (23) | 8 (50) | 3 (27) | 1 (33) | 18 (32) |
| **<u>Pertinent Medical and Physical Conditions</u>** |  |  |  |  |  |
| Family history (Yes) – n (%) | 19 (73) <sup>a</sup> | 2 (12) | 3 (21) <sup>c</sup> | 3 (100) | 27 (48) |
| Chronic pain in body (Yes) – n (%) | 8 (31) | 9 (56) | 3 (27) | 1 (33) | 21 (37) |
| Bisphosphonate (Yes) – n (%) | 11 (42) | 16 (100) | 11 (100) | 3 (100) | 41 (73) |
| Wheelchair use (Yes) – n (%) | 1 (4) <sup>a</sup> | 14 (88) <sup>b</sup> | 4 (36) <sup>c</sup> | 2 (67) | 21 (38) |
| Days missed school – mean (SD) | 9 (12) <sup>a</sup> | 25 (24) | 16 (14) <sup>c</sup> | 29 (29) | 16 (18) |

**Statistical tests**) determine the significant relationship between categorical variables and OI types I, III and IV: Chi-square test or the Fisher's exact test for contingency tables with small cell counts; Compare means of a continuous variable between OI types I, III and IV: Welch's t-test for independent samples. As the sample size is small in each group (n<15), results have been confirmed by Mann-Whitney U test (non-parametric test).
<sup>a</sup> p<0.05 OI type I compared to OI type III.
<sup>b</sup> p<0.05 OI type III compared to OI type IV.
<sup>c</sup> p<0.05 OI type IV compared to OI type I.

**Table 2.** The Child Perceptions Questionnaire subscales for 8 to 10-year-old children (CPQ8–10)

| OHRQoL | Number of items | Possible range | Observed range | OI I<br>(n=26) | OI III<br>(n=16) | OI IV<br>(n=11) | Others<br>(n=3) | Total<br>(n=56) |
| --- | --- | --- | --- | --- | --- | --- | --- | --- |
| Overall | 25 | 0-100 | 0-43 | 10.0 (10.5) | 9.8 (6.4) | 9.0 (7.3) | 9 (6.2) | 9.7 (8.5) |
| Oral symptoms | 5 | 0-20 | 0-15 | 4.9 (3.8) | 4.8 (2.6) | 4.4 (1.5) | 5.3 (3.1) | 4.8 (3.1) |
| Functional Limitation | 5 | 0-20 | 0-8 | 1.3 (1.9) | 2.6 (2.5) | 1.4 (2.2) | 2.0 (1.7) | 1.7 (2.2) |
| Emotional Well-Being | 5 | 0-20 | 0-20 | 1.9 (4.2) | 1.7 (2.3) | 1.7 (3.8) | 0.6 (1.2) | 1.8 (3.5) |
| Social Well-Being | 10 | 0-40 | 0-13 | 1.8 (3.3) | 0.7 (1.3) | 1.5 (2.7) | 1.0 (1.0) | 1.4 (2.7) |

Results are shown as n or mean (SD).
**Statistical analysis:** Welch's t-test, results have been confirmed by Mann-Whitney U test (non-parametric test).
<sup>a</sup> $p < 0.05$ OI type I compared to OI types III.
<sup>b</sup> $p < 0.05$ OI type III compared to OI types IV.
<sup>c</sup> $p < 0.05$ OI type IV compared to OI types I.

In adolescents aged 11 to 14 years (Table 3), more teenagers with OI types I were having a parent living with OI compared with OI type III (p-value <0.05). Having chronic pain throughout the body was more prominent in OI type III compared to OI type I (p-value <0.05). All individuals with OI type III and most with OI type IV had a history of bisphosphonate treatment (oral or IV), compared with 51% in OI type I. Using a wheelchair as a mean of transportation was more prevalent in OI type III compared to OI types I and IV (p-value <0.05) and the number of wheelchair users were significantly higher in OI type IV. The number of days that teens missed school related to OI was significantly higher in type III compared to type I (p-value <0.05) and also significantly higher in OI type IV compare to OI type I (p-value <0.05). Total scores of the CPQ11–14 were significantly higher (worse) in OI types III or IV compared to type I (p-value <0.05 for both). When the sub-scales were compared, functional limitations had a greater negative impact on the OHRQoL of adolescents suffering from OI type III or IV (p-value <0.05 for both) when compared to those suffering from OI type I (Table 4).

**Table 3.** Characteristics of the 11 to 14-year-old group

| Patients aged between 10-14 | OI I | OI III | OI IV | Others | All |
| --- | --- | --- | --- | --- | --- |
| **<u>Sociodemographic Characteristics</u>** |  |  |  |  |  |
| Enrolment number – n (%) | 39 (48) | 14 (17) | 23 (28) | 6 (7) | 82 (100) |
| Female | 22 (56) | 11 (79) | 14 (61) | 4 (67) | 51 (62) |
| Age – mean (SD) | 13.2 (1.3) | 13.4 (1.1) | 13.1 (1.2) | 13.7 (1.2) | 13.2 (1.2) |
| Race (White) – n (%) | 32 (82) | 12 (86) | 19 (83) | 4 (67) | 67 (82) |
| others | 7 (18) | 2 (14) | 4 (17) | 2 (33) | 15 (18) |
| Insurance status (Private) – n (%) | 26 (67) | 9 (64) | 14 (61) | 3 (50) | 52 (63) |
| Medicare/Medicaid | 13 (33) | 5 (36) | 9 (39) | 3 (50) | 30 (37) |
| **<u>Pertinent Medical and Physical Conditions</u>** |  |  |  |  |  |
| Family history (Yes) – n (%) | 23 (59) <sup>a</sup> | 1 (7) | 7 (30) | 3 (50) | 34 (41) |
| Chronic pain in body (Yes) – n (%) | 11 (28) <sup>a</sup> | 10 (71) | 9 (39) | 4 (67) | 34 (41) |
| Bisphosphonate (Yes) – n (%) | 20 (51) | 14 (100) | 21 (91) | 4 (67) | 59 (72) |
| Wheelchair use (Yes) – n (%) | 1 (3) <sup>a</sup> | 13 (93) <sup>b</sup> | 10 (43) <sup>c</sup> | 4 (67) | 28 (34) |
| Days missed school – mean (SD) | 9.2 (10.4) <sup>a</sup> | 17.1 (15.6) | 27.2 (21.8) <sup>c</sup> | 14.7 (13.8) | 16.2 (17.1) |

**Statistical tests)** determine the significant relationship between categorical variables and OI types I, III and IV: Chi-square test or the Fisher's exact test for contingency tables with small cell counts; Compare means of a continuous variable between OI types I, III and IV: Welch's t-test for independent samples. As the sample size is small in each group (n<15), results have been confirmed by Mann-Whitney U test (non-parametric test).
<sup>a</sup> p<0.05 OI type I compared to OI types III.
<sup>b</sup> p<0.05 OI type III compared to OI types IV.
<sup>c</sup> p<0.05 OI type IV compared to OI types I.

**Table 4.** The Child Perceptions Questionnaire subscales for 11 to 14-year-old children (CPQ11-14)

| OHRQoL | Number of items | Possible range | Observed range | OI I<br>(n=39) | OI III<br>(n=14) | OI IV<br>(n=23) | Others<br>(n=6) | Total<br>(n=82) |
| --- | --- | --- | --- | --- | --- | --- | --- | --- |
| Overall | 37 | 0-148 | 1-53 | 16.5 (12.8)<sup>a</sup> | 24.7 (12.5) | 23.1 (14.4)<sup>c</sup> | 22.3 (17.7) | 20.2 (13.8) |
| Oral symptoms | 6 | 0-24 | 1-11 | 5.8 (2.9) | 7.1 (3.2) | 7.1 (3.2) | 6.7 (4.3) | 6.4 (3.1) |
| Functional Limitation | 9 | 0-36 | 0-19 | 4.3 (4.2)<sup>a</sup> | 8.6 (5.1) | 7.2 (4.9)<sup>c</sup> | 7.4 (5.9) | 6.1 (4.9) |
| Emotional Well-Being | 9 | 0-36 | 0-20 | 3.6 (5.7) | 5.7 (5.9) | 5.5 (6.9) | 5.3 (7.2) | 4.6 (6.2) |
| Social Well-Being | 13 | 0-52 | 0-19 | 2.9 (4.7) | 3.3 (3.8) | 3.5 (4.5) | 3.0 (3.1) | 3.1 (4.3) |

Results are shown as n or mean (SD).
Statistical analysis: Welch's t-test, results have been confirmed by Mann-Whitney U test (non-parametric test).
<sup>a</sup> p<0.05 OI type I compared to OI types III.
<sup>b</sup> p<0.05 OI type III compared to OI types IV.
<sup>c</sup> p<0.05 OI type IV compared to OI types I.

Tables 5 and 6 show the results of multivariable-adjusted ordinal logistic regression for children and teenagers (respectively). Among children, a diagnosis with the more severe type of OI (type IV and III, respectively) was not associated with a negative impact on OHRQoL. Although not statistically significant but having OI type III or IV among children were associated (p>0.05) with a higher grade of functional limitations domain compared to type I (Table 5). However, this association was statistically significant amongst teenagers (Table 6). After adjusting for sociodemographic variables and family history of OI, adolescents having OI type III compared to OI type I were more likely to have a higher (worse) score of CPQ_11-14_ (P<0.05). This association was predominantly attributed to the strong correlation between OI types and functional limitations domain (subscale of CPQ). Although the total CPQ_11-14_ score for OI type IV was significantly higher than for OI type I in univariate analysis, this difference became statistically insignificant after adjusting for the other variables in the model. However, the difference between OI types IV and I persisted with regard to functional limitation (P<0.05). Having OI type III among adolescents was statistically significantly (P<0.05) associated with a higher grade of functional limitations domain compared to OI type I.

**Table 5.**
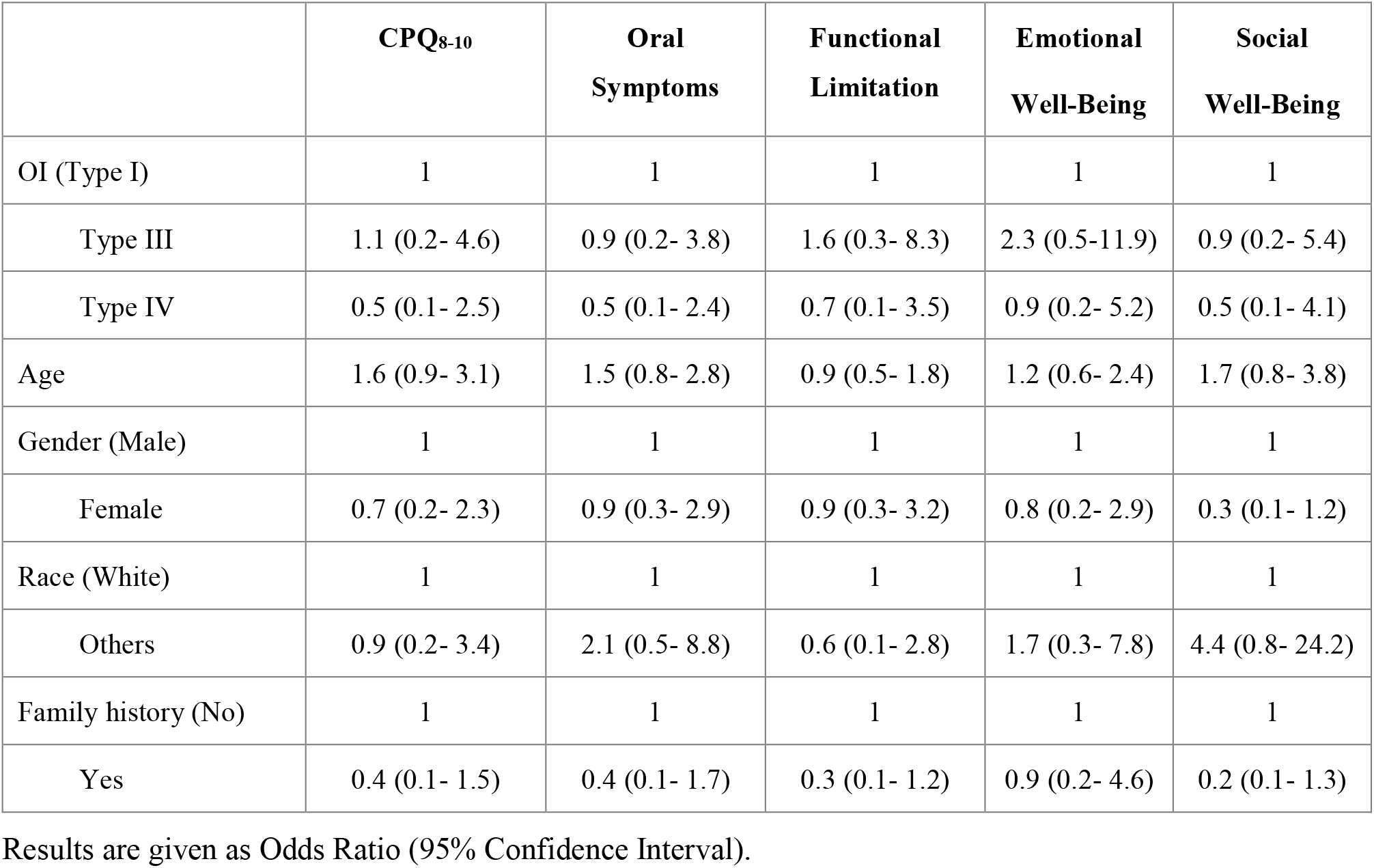
Adjusted odds ratio of the negative impact (having higher grades) on the oral health-related quality of life in children aged 8- 10 years

**Table 6.** Adjusted odds ratio of the negative impact (having higher grades) on the oral health-related quality of life in adolescents aged 11 - 14 years

|  | **CPQ<sub>11-14</sub>** | **Oral**<br>**Symptoms** | **Functional**<br>**Limitation** | **Emotional**<br>**Well-Being** | **Social**<br>**Well-Being** |
| --- | --- | --- | --- | --- | --- |
| OI (Type I) | 1 | 1 | 1 | 1 | 1 |
| Type III | 4.6 (1.2- 17.4) * | 1.8 (0.5- 6.6) | 7.8 (1.9- 31.7) * | 3.6 (0.9- 13.7) | 0.9 (0.3- 3.6) |
| Type IV | 2.6 (0.9- 7.3) | 1.7 (0.6- 4.6) | 3.7 (1.3- 10.3) * | 2.3 (0.8- 6.6) | 1.7 (0.7- 4.7) |
| Age (years) | 1.1 (0.8- 1.6) | 1.1 (0.8- 1.6) | 1.2 (0.8- 1.6) | 1.1 (0.7- 1.6) | 0.9 (0.6- 1.3) |
| Gender (Male) | 1 | 1 | 1 | 1 | 1 |
| Female | 0.9 (0.4- 2.5) | 0.8 (0.3- 2.1) | 0.7 (0.3- 1.9) | 0.8 (0.3- 2.1) | 1.6 (0.7- 3.9) |
| Race (White) | 1 | 1 | 1 | 1 | 1 |
| Others | 2.3 (0.6- 8.3) | 1.8 (0.6- 6.3) | 1.2 (0.4- 4.2) | 2.44 (0.7- 8.9) | 1.4 (0.4- 4.5) |
| Family history (No) | 1 | 1 | 1 | 1 | 1 |
| Yes | 1.1 (0.4- 2.6) | 1.1 (0.4- 2.9) | 0.9 (0.4- 2.6) | 1.3 (0.5- 3.4) | 0.9 (0.3- 2.2) |

Results are given as Odds Ratio (95% Confidence Interval).
\* Statistically Significant findings at $p < 0.05$

## Discussion

In this study we found that adolescents with OI type III had a more negative overall profile of OHRQoL when compared to OI type I. Functional limitation seems to negatively affect OHRQoL of OI types III and IV in comparison with type I amongst teenagers. While sharing similar levels of OHRQoL in three domains (oral symptoms, emotional wellbeing, and social wellbeing), teens with moderate and severe OI (types IV and III) reported worse functional OHRQoL compared to mild OI (type I). This result shows that although the functional limitations in moderate and severe OI affects their perception of physical (functional) oral health QoL, it does not influence their perception of mental and psychological oral health QoL when compared with mild OI.

In general, our results in children and teenagers with OI show a better OHRQoL than what has been observed in populations of the same age with a variety of conditions [10, 11, 21]. In children with common dental disease (caries) and orofacial conditions (lip and palate cleft) mean CPQ8-10 scores of 19.1 and 18.4 were found [10], compared to a mean CPQ8-10 score of 9.7 in the present study. This suggests a better OHRQoL amongst children with OI.

Among 11 to 14-year-old adolescents, we found mean CPQ11-14 scores of 24.7 and 23.1 for OI types III and IV, respectively, which is similar to what has been reported for otherwise healthy adolescents with orthodontic disorders and dental caries but better than in adolescents with orofacial conditions (mean CPQ11-14 score: 31.4) [11]. Adolescents with OI type I had significantly lower CPQ11-14 scores compared to the other OI groups or the cohorts reported in the literature.

One interesting observation of the present study was that OHRQoL did not vary between OI types in 8 to 10 year-old children, but was significantly worse in 11 to 14 year-old adolescents with severe OI than in adolescents with mild OI. One explanation for this discrepancy between age groups is that the adolescents experienced functional problems over a longer period of time or perhaps are more conscious of the difficulties caused by the disease.

Quality of life is not static; it is a complex, multifaceted, and dynamic construct that can be defined and measured. It varies between individuals and it changes within the same individual overtime. Consequently, the results of any QoL evaluation has an inherent instability. The relation between oral health status (symptoms) and QoL is not simple nor is it a direct relationship. “People assess their HRQoL by comparing their expectations and experiences” [21]. Expectations are altered and learned by and from experiences. Questionnaires used to evaluate OHRQoL measures the gap between the expectations (hopes) of the individual, and it’s present experience. Existing measures of OHRQoL do not account for expectations of oral health; they only detect the negative impact of the disease or treatment on patient’s perception of oral health QoL. Moreover, patients may be at different points of their illness trajectory when their quality of life is measured [21, 22]. With this understanding, one can interpret the results of this study that the discrepancy (gap) between expectation level and individuals experience is significantly higher in OI types III compare to OI type I at the specific time that their OHRQoL data has been collected with functional limitation being the main contributor. This difference may be due to higher level of expectations or more deteriorating experiences (severe symptoms) or a combination thereof in OI types III compared to those conflicted with OI type I. Therefore, types III are more vulnerable to have lower score in OHRQoL compared to OI type I. (Figure 1)

**Figure 1.**
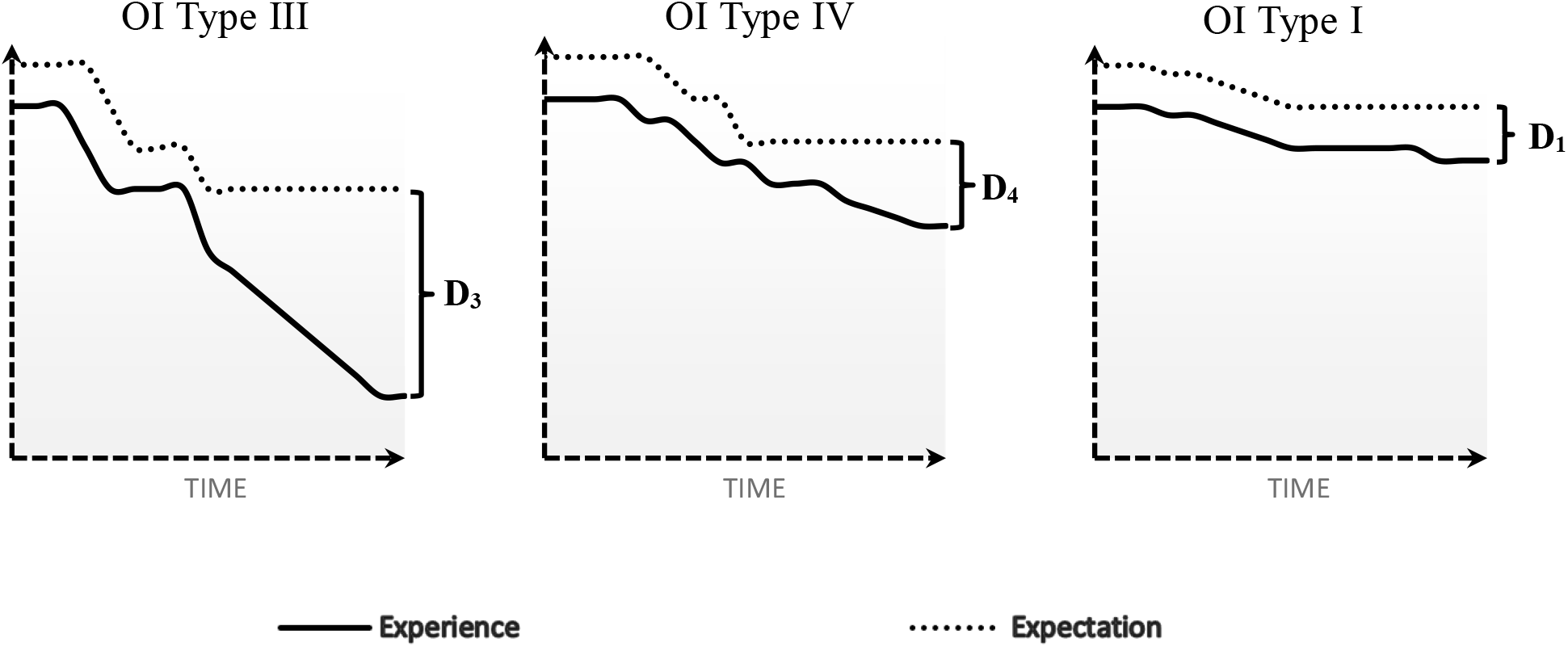
Schematic diagram of the differences in expectation-experience between OI types **(D3>D1; p<0.05)**

Physicians, dentists, and caregivers, in an attempt to provide good care, are trying to bridge the gap between the patients’ experiences and their expectations. In other words, how patients want to be and what their physical health allows them to be. This is commonly achieved through clinical interventions to restore impairments (improve experience) or administering psychological interventions to help them adjust their expectations to their altered clinical health status (diminish expectation).

The OHRQoL scores obtained in this study employing CPQ questionnaire show that these instruments are likely to have “floor effects”, which signifies that they cannot demonstrate improvement in their postinterventional condition. Therefore, given the unique dental issues in OI, it may be useful to develop a more tailored measure for assessing OHRQoL for this population. Disease-specific instruments are generally more sensitive to oral health traits of this condition and more responsive to changes during the time (less “floor effect”) in comparison with generic measures [23-24].

## Conclusions

In conclusion, this study found that teens with OI type III and IV have higher grades of functional limitations than OI type I. This association leads to a lower OHRQoL in teens with OI type III compared with OI type I.

## Abbreviations

OI: Osteogenesis Imperfecta
OHRQoL: Oral Health-Related Quality of Life
DI: Dentinogenesis Imperfecta
WHO: World Health Organization
QoL: Quality of Life
HRQoL: Health-Related Quality of Life
CPQ: Child Perceptions Questionnaire

## Ethics approval and consent to participate

The study obtained ethics approval from McGill ethics committee, number A09-M47-15B, and all study participants or their legal guardians provided informed consent.

## Consent for publication

Not applicable

## Availability of data and material

The datasets used and/or analyzed during the current study are available from the corresponding author on reasonable request.

## Authors’ contributions

Performed data-cleaning, statistical analysis, and wrote the manuscript: MN. Conceived and designed the experiment, obtained ethical approvals, and securing research fundings: FR, VRS, BL, J-MR. Identified and recruited OI patients: J-MR. Assissted in drafting the manuscript as well as revising it critically for important intellectual content: SE, FR, MSM.

## Acknowledgments

We gratefully acknowledge the contributions of members of the BBD (Brittle Bone Disease) Consortium namely: Sandesh CS Nagamani, Francis Glorieux, Paul Esposito, Eric Rush, Michael Bober, David Eyre, Danielle Gomez, Gerald Harris, Tracy Hart, Mahim Jain, Deborah Krakow, Jeffrey Krischer, Eric Orwoll, Cathleen Raggio, Peter Smith, and Laura Tosi.

